# The extremely low mechanical force generated by nano-pulling induces global changes in the microtubule network, nuclear morphology, and chromatin transcription in neurons

**DOI:** 10.1101/2025.03.09.641873

**Authors:** Alessandro Falconieri, Lorenzo Da Palmata, Valentina Cappello, Tiziana Julia Nadjeschda Schmidt, Pietro Folino, Barbara Storti, Ranieri Bizzarri, Vittoria Raffa

## Abstract

Mechanical force plays a pivotal role in every aspect of axon development. In this paper, we explore the use of nano-pulling, a technology that enables the intracellular generation of extremely low mechanical forces. We demonstrate that force-mediated axon growth also exerts global effects that extend to the nuclear level. Our mechanistic studies support a model in which exogenous forces induce stabilization of microtubules, and a significant remodeling of perinuclear microtubules, which preferentially align perpendicularly to the nuclear envelope. We observed an increase in the lateral tension of the nucleus, leading to substantial remodelling of nuclear morphology, characterized by an increase in nuclear grooves and higher sphericity index (indicating less flattened nuclei). Notably, these changes in nuclear shape are linked to chromatin remodelling, resulting in global transcriptional activation.

## 1 Introduction

Neurons, like other cell types, are continuously exposed to mechanical and topological cues. To adapt or respond to these stimuli, they have evolved several mechanotransduction pathways.[1,2] One of the most well-characterized processes in this context is axon navigation in response to mechanical forces. The role of force as a driver of axonal elongation was first identified in 1941 when Paul Weiss coined the term “towed growth” to describe the mechanism by which neurons are physically drawn out as the body grows.[3] In 2004, Smith et al. introduced the term “stretch-growth”, referring to the force-induced axonal outgrowth.[4] Today, mechanical force is recognized as a critical extrinsic factor in modulating various axonal processes, including axon pathfinding, elongation, initiation, pruning, fasciculation, and synapse formation.[5] These mechanisms are typically localized in the peripheral regions of neurons, such as axons and dendrites. However, research over the past decade on mechanotransduction across various models—ranging from prokaryotes,[6] yeasts,[7] plants,[8] and mammalian cells[9] —suggests that mechanotransduction should be understood as a global process. Upon sensing mechanical stimuli, a fast remodelling of the cell occurs within minutes to hours. This reorganization efficiently propagates the response across multiple cellular compartments, leading to a global adaptation.[10] In neurons, however, this topic remains largely under-explored. In this study, we aim to investigate whether the application of exogenous force that promotes axon growth can also influence other compartments, particularly the nucleus. In cells, the cytoskeleton is mechanically connected to the nucleoskeleton through the linker of nucleoskeleton and cytoskeleton (LINC) complex proteins.[11,12] Through the LINC complex, the outer nuclear membrane (ONM) binds to the contractile cytoskeleton,[13–15] and the inner nuclear membrane (INM) recruits nuclear lamins.[16] Lamins, which are intermediate filaments with load-bearing mechanical properties,[17] buffer mechanical stimuli originating both inside and outside the nucleus. Lamins tether heterochromatin to the INM at the lamin-associated chromatin domains.[18] Thanks to this connection, lamins can dissipate mechanical stress by altering the global 3D chromatin network, inducing subnuclear movements and chromatin mobility.[19]

The physical link between cytoskeleton and chromatin fibers raises the possibility that exogenous force applied to neurons can induce a remodelling at the nuclear level.[20] However, when studying cell mechanotransduction, it is crucial to consider the type of mechanical stimulation, as mechanical force can vary in type of load (e.g., traction, shear stress, or compressive stress), and the resulting cellular responses may be dependent on time (acute or chronic stimulus), load (distributed or point force), direction and magnitude of the force vector.[21] Many cell types are mechanosensitive, yet they do not all respond to the same force range. Cardiomyocytes, smooth muscle cells, osteoblasts, and glial cells typically respond to nanoNewton (nN) but not to picoNewton (pN) forces. For instance, the application of a 20% strain (corresponding to tens of nN) induces a rearrangement of the glial cell network, affecting the distance and orientation between nuclei,[22] while axons break and disconnect within seconds under a strain above 2% (equivalent to a few nN).[4] Conversely, pN forces applied to axons influence all stages of axonal development.[5,23,24] Based on this rationale, in this study, we employed magnetic nano-pulling, a technology capable of generating forces in the pN range.[25] Our findings indicate that developing axons of neurons subjected to chronic (days) stretching via magnetic nano-pulling nearly double their elongation rate and halve the time required for synaptic maturation.[25] This technology simulates the chronic exposure to mechanical forces that neurons experience during physiological processes, such as the strain that spinal neurons undergo during body growth, which occurs at approximately 5 μm/h, the same rate induced by nano-pulling.[25] By applying these chronic, extremely low forces to hippocampal neurons, we examined the nuclear-level response and sought to elucidate the underlying mechanisms. Our data support the hypothesis that force induces a remodelling of the microtubule network, alterations in nuclear shape and ultimately global chromatin reorganization.

## 2 Results

### 2.1 Force induces MT stabilization and re-organization of the MT network in the soma

Nano-pulling is a methodology that consists of loading axons with superparamagnetic iron oxide nanoparticles (SPIONs) to remotely generate a precise force vector through external magnetic fields (Figure 1A). In previous studies, we estimated the generated force in the order of 10 pN [25,26]. This force was found to increase axon length from 140±4 µm to 288±7 µm (n=120 axons, p<0.0001) without reducing the caliber of the axon (4.8±0.2 µm for control axons and 5.2±0.2 µm for stretched axons, n=24 axons, p=0.1). This corresponds to an increase in elongation rate from approximately 2 to 5 µm/h in stretch-growth compared to spontaneous elongation (tip-growth). In a previous publication, we demonstrated that the nano-pulling leads to stabilization of axonal MTs that, in turn, induces an accumulation of MTs in the axon shaft[27]. Specifically, axons stretched for 48 h or 120 h show a similar increase (≈40%) in the number of axonal microtubules compared to control axons [27]. Since α-tubulin acetylation is associated with stable MTs,[28] whereas tyrosination correlates with dynamic instability, [29] the ratio between α-tubulin acetylation and tyrosination levels is usually used as a marker of MT stability.[27,30,31] In this study, we evaluated whether nano-pulling induces MT stabilization in the soma, as well. Analysis of ac/tyr ratio of α-tubulin in the axonal compartment confirmed the force-mediated MT stabilization (Figure 1C1, 2.18±0.08 for control axons and 2.43±0.09 for stretched axons, n=60, p=0.035), in analogy with our previous observations [27,31]. The analysis in the soma showed a value of 1.08±0.03 for control neurons, which was significantly different from the value for stretched neurons (1.29±0.04) (Figure 1C2, n=75, p=0.002), suggesting that force-mediated stabilization occurs also in the soma.

**Figure 1.**
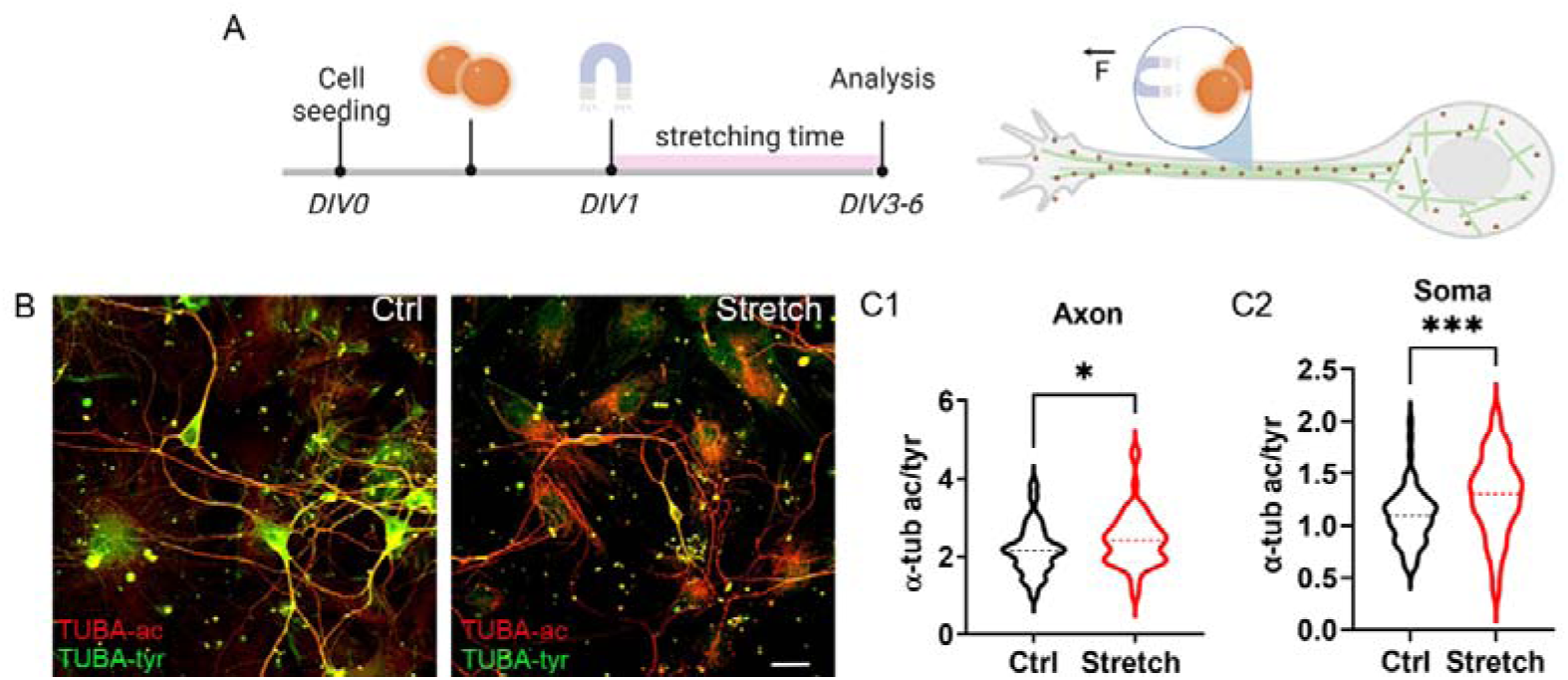
Nano-pulling induced MT stabilization. A) Loading protocol of hippocampal neurons with SPIONs. SPIONs are added in cell growth media at the concentration of 5 µg/ml four hours after cell seeding (day in vitro, DIV0) and the magnet is applied from DIV1. Neurons are stretched for the subsequent 48 h (if not indicated differently). B) Immunofluorescence against acetylated α-tubulin (red) and tyrosinated α-tubulin (green) in control and stretched neurons. Scale bars: 20 µm. C1) Ratio of acetylated versus tyrosinated α-tubulin in axons. Violin plot, n=60 axons from four replicates, unpaired t test, 2-tailed, p=0.035, t=2.131 df=118. C2) Ratio of acetylated versus tyrosinated α-tubulin in neuron soma. Violin plot, n=75 neurons from four replicates, unpaired t test, 2-tailed, p=0.0002, t=3.807, df=148.

Next, we focused the analysis on soma to identify changes in the MT network following the increase in MT stability. Cells were formalin-fixed and the ultrastructure was studied via TEM. MT orientation was studied in the region proximal to the outer side of the nuclear envelope. In the soma of control neurons, the MTs near the nuclear lamina grow preferentially parallel to it. Figure 2A1 shows a representative image of a control neuron where MTs close to the nuclear envelope present a preferential tangential orientation that can be parallel (red lines) or perpendicular to the cutting plane (red spot). Conversely, stretched neurons show many MTs that are mostly oriented orthogonal to the nuclear envelope (Figure 2A2). By analysing more than seventy neurons, randomly selected in both groups, we found that the percentage of neurons showing MTs with orientation orthogonal to the nuclear membrane is very low in controls (11%) but significantly higher in stretched group (72%) (Figure 2B2). To investigate this further, we analyzed the morphology of the nuclear envelope. Nuclear compartments appear clearly detectable in the soma of neurons arising from both the experimental classes. Nucleoplasm and cytoplasm are divided by a well-defined nuclear envelope. Nuclear pores can be observed along the nuclear membrane (inset a of Figure2A1, green rectangle, scale bar 200 nm). When the envelope is tangential to the cutting plane, the nuclear pores appear as small circular structures (inset b of Figure2A1, green circle, scale bar 200 nm). It is also possible to observe the accumulation of peripheral heterochromatin that forms regions of densely packed DNA in the proximity of the inner side of the nuclear envelope. Interestingly, the presence of MTs oriented orthogonally to the envelope appears to correlate with the presence of grooves (invaginations/protrusions) in the nuclear envelope (Figure 2A2/A3). We observed an increased amount of grooves (invaginations/protrusion) of the nuclear envelope in the stretched group (Figure 2A2/A3) compared with the control group (Figure 2A1). As a consequence of the increase in nuclear envelope roughness, more nuclear pores are detectable in the stretched group. In fact, when the nuclear shape is characterised by many invaginations/protrusions, both parallel and orthogonal to the cutting plane, it presents a higher number of tangential surfaces in which it is possible to observe nuclear pores (Figure 2A3, green spots). The presence of grooves was quantified with the parameter α, defined as the ratio between the perimeters of the nuclear envelope and its convex hull (see methods). Experimental data confirm that nuclei of stretched neurons have a more jagged contour (Figure 2B3, p<0.0001).

**Figure 2.**
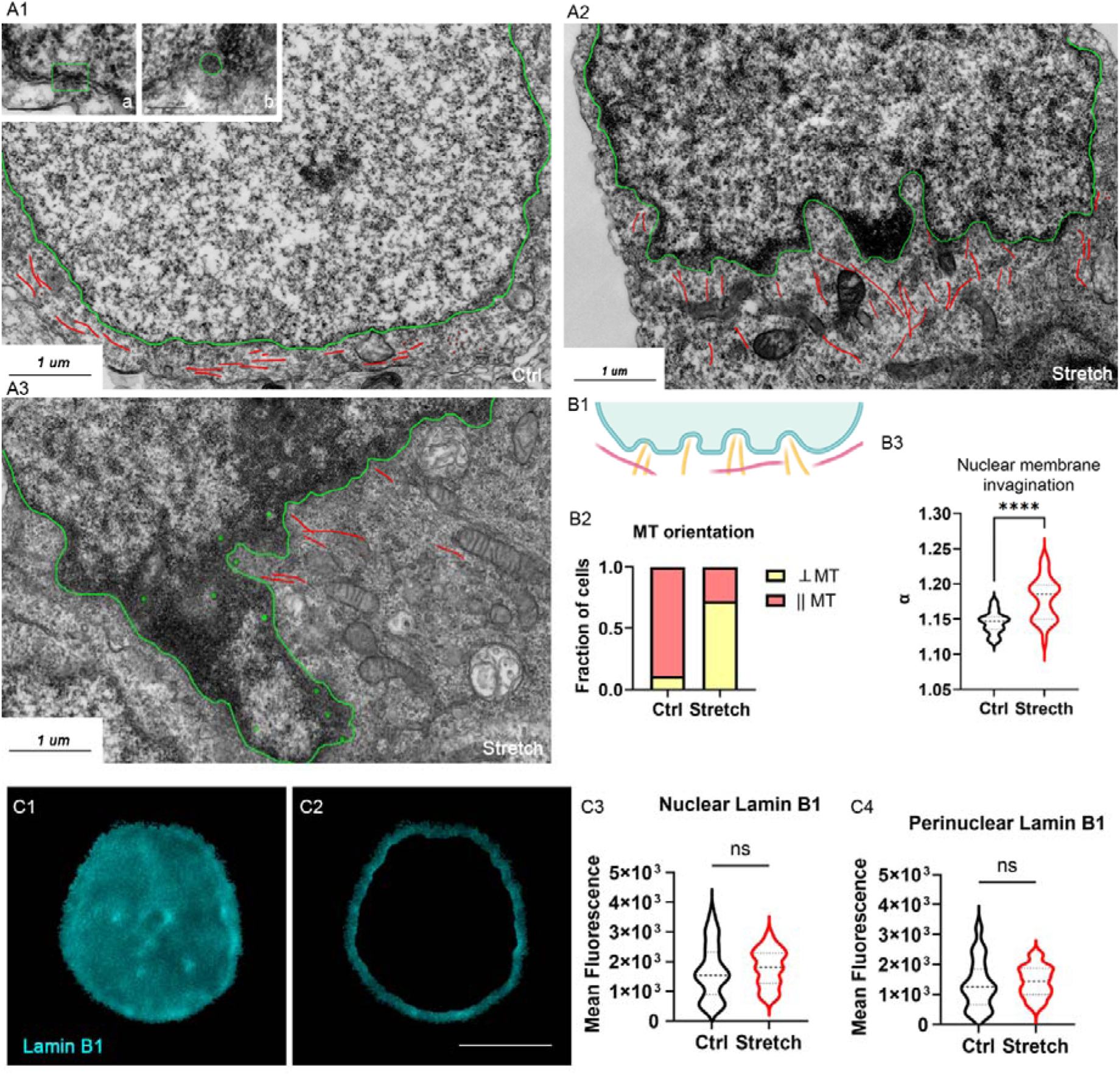
Perinuclear cytoskeleton re-organization. A) TEM images of cross-sections of control (A1) and stretched neurons (A2, A3). Red lines and dots show MTs that are longitudinal or orthogonal to the nuclear envelope, respectively. Green lines mark the nuclear envelope, green dots mark nuclear pores. Inset a and b of panel A1 show a longitudinal and transversal cross section of nuclear pores, respectively (scale bar, 200 nm). B1) Scheme of nuclear envelope invagination. MT perpendicular to the envelope shown in red, the MT parallel to the envelope in yellow. B2) Fraction of cells presenting MT perpendicular to the envelope from n=75 neurons from two replicates. Chi-square test performed on integer numbers, Fisher’s exact test, p<0.0001. B3) α value (ratio between the perimeter of nuclear envelope and the perimeter on the convex hull) between control and stretched neurons (violin plot). Unpaired t test, two-tailed, n=25 nuclei from two replicates, t=4,942, df=44, p <0.0001. C1) Lamin B1 immunostaining in the nucleus. C2) Lamin B1 immunostaining, in a perinuclear ring of 500 nm. Scale bar: 5 μm. C3) Mean fluorescence of the nucleus. Unpaired t test, n=60 from four replicates, p=0.45, t=0.7620, df=117. C4) Mean fluorescence of the perinuclear ring. Mann Whitney test, n=60, p=0.23, Mann-Whitney U=1543.

Since the loss of lamin B1 was recently associated with nuclear deformations in hippocampal neurons,[32] after immunostaining for lamin B1, nuclei were observed using super-resolution fluorescence microscopy. We quantified the total and mean fluorescence in the nucleus (Figure 2C1) and in a perinuclear ring of 500 nm thickness (Figure 2C2). We did not observe any change in nuclear mean fluorescence (1.69±0.11·10³ and 1.80±0.08·10³ in control and stretched nuclei, respectively, p=0.45, Figure 2C3), total fluorescence (integrated density: 1.66±0.81·10C and 1.40±0.42·10C in control and stretched nuclei, respectively, p=0.45, data not shown), or mean fluorescence in the perinuclear region (1.35±0.11·10³ and 1.44±0.07·10³ in control and stretched nuclei, respectively, p=0.23, Figure 2C4), indicating no detectable changes in lamin B1 between the two conditions.

### 2.2 Force induces a remodelling of nuclear shape that is dependent on cytoskeleton

Microtubules physically interact with the nuclear membrane. The forces generated by cytoplasmic microtubules alter and control nuclear shape [33,34]. Here, we were wondering whether the re-organization of perinuclear MTs in stretched cells, oriented perpendicularly to the nuclear envelope, could induce changes in nuclear morphology. A change in nuclear shape seems to be suggested by TEM analysis, documenting a variation of the circularity value (min Feret/max Feret) between the two conditions (0.63±0.18 and 0.51±0.19, for control and stretched nuclei, respectively; n=76; Mann Whitney test; p= 0.0002). However, TEM imaging does not allow for the reconstruction of 3D nuclear morphology because the analysis is based on cell pellets and the cutting plane is random. Indeed, as next step, we analysed cell monolayers by super-resolution confocal microscopy, we reconstructed the 3D morphology by the plugin 3D Ellipsoid - 3D Suite ImageJ[35], and we derived the morphological parameters, i.e., the nuclear volume *V*, the area *S* (Z projection of the volume), the major axis *d* and the heigh *h* calculated from the interpolated ellipse (geometrical parameters shown in Figure 3A). The nuclear volume was not different between the two conditions (Figure 3C1, p=0.87), suggesting that morphological changes are likely due to the elastic deformation of the nuclear shape rather than a change in nuclear size. We found that the nano-pulling reduced the length of the major axis *d* (7.5±0.1 and 6.5±0.1 µm in control and stretched neurons, respectively, p<0.0001, Figure 3C2), increase the ellipse height *h* (2.73±0.05 and 3.25±0.07 µm in control and stretched neurons, respectively, p<0.0001, Figure 3C3) and increase sphericity (0.342±0.008 and 0.389±0.009 in control and stretched neurons, respectively, p=0.0002, Figure 3C4). The area *S* was statistically significant smaller in stretched neurons compared to control neurons (60.08±1.27 μm^2^ and 49.15±1.13 μm^2^ in control and stretched neurons, respectively, Figure 3D1, no pharmacological treatment). These changes in the morphological parameters *S*, *d*, and *h* are consistent with a more laterally compressed nucleus in the stretched neurons compared to the control neurons. Figure 3B shows representative images generated from a 3D reconstruction based on a z-stack, illustrating the net change in nuclear morphology in stretched neurons. Since growing protofilaments generate a pushing force,[36] we wondered whether this nuclear compression could be driven by MTs, which are perpendicularly oriented to the nuclear envelope in stretched neurons but not in control neurons. To investigate this further, we measured the *S* parameter when cells were treated with Nocodazole (Ncz, 1.8 ng/ml), a drug that interfere with MT polymerization.[37] This concentration has no effect on control cells (p=0.67 for Ncz+ vs Ncz-control groups) but totally blocked the change in nuclear shape induced by the nano-pulling (in Ncz treated groups, S was 58.08±1.13 μm^2^ and 58.88±1.45 μm^2^ in control and stretched neurons, respectively, p=1, Figure 3D1).

**Figure 3.**
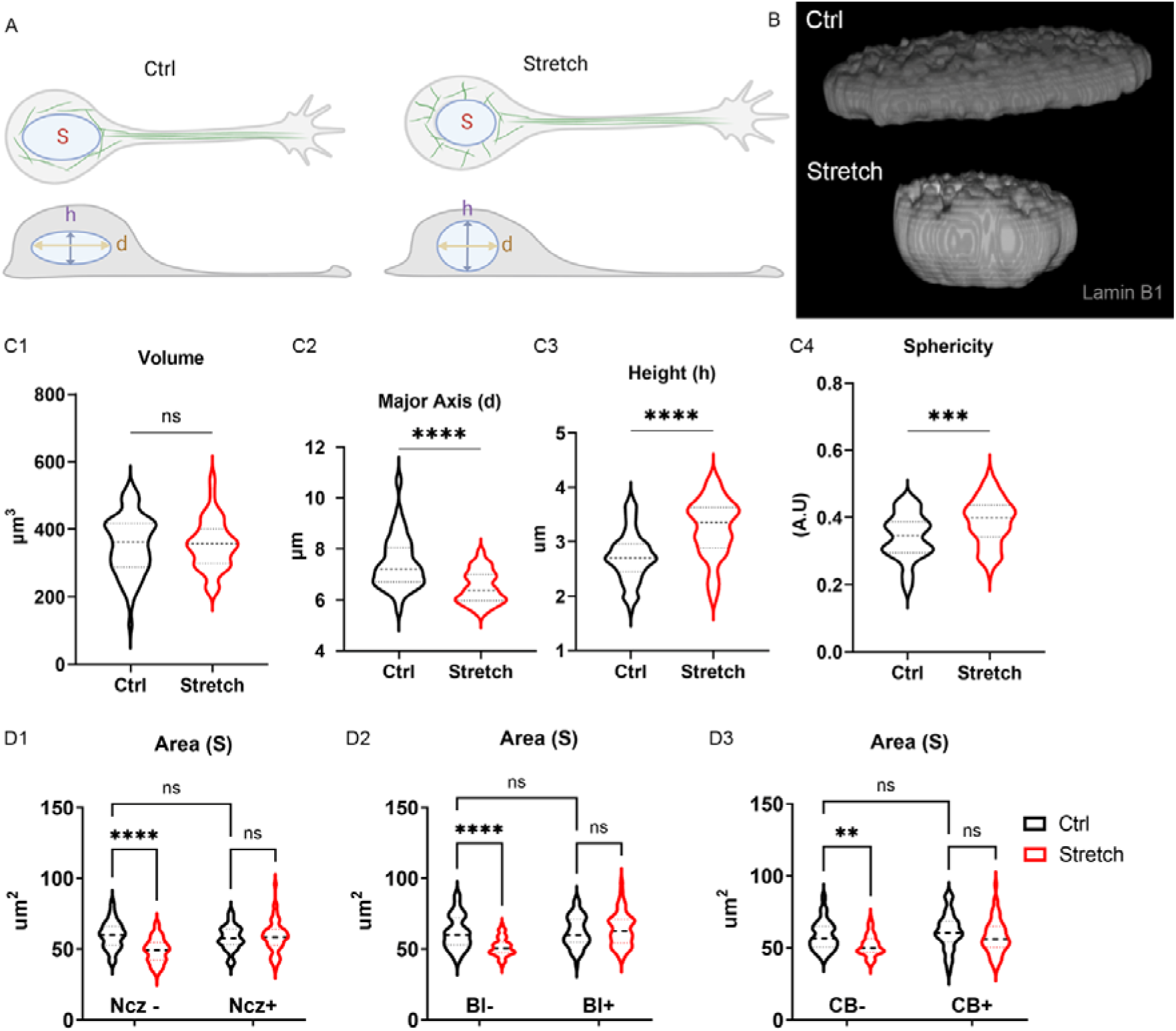
Nuclei remodelling. A) The scheme shows the different organization of MT (green lines) in control and stretched neuron and the nuclear morphological parameters: 2D projected area (S), major axis (d) and height from the basal plane (h). B) Representative images of nuclei of control neurons (flatter) and stretched neurons (more spherical). Lamin B1 staining. (3D rendering is available in supplementary video). C1) Nuclear volume (violin plot). Unpaired t-test, p=0.96, t=0.04634, df=127. n=60 neurons from four replicates. C2) Nuclear major axis (violin plot). Mann-Whitney test, p<0.0001. n=60 neurons from four replicates. C3) Nuclear height (violin plot). Mann-Whitney test, p<0.0001. n=60 neurons from four replicates. C3) Nuclear sphericity (violin plot). Unpaired t-test, p=0.0002, t=3,818, df=126. n=60 neurons from four assays. D) Analysis of the nuclear cross-section area (S). D1) Nocodazole was added at DIV1 and S value measured at DIV3 (violin plot). 2-way ANOVA test, with post-hoc Tukey HSD. Row factor (Ncz-/+): p=0.0023, F=9.503. Columns factor (Ctrl/Stretch): p<0.0001, F=16.34. n=60 neurons from four replicates. D2) 0.4 ug/ml Blebbistatin was added at DIV1 and S value measured at DIV3 (violin plot). 2-way ANOVA test, with post-hoc Tukey HSD. Row factor (Bl-/+): p<0.0001, F=22.56. Columns factor (Ctrl/Stretch): p=0.0021, F=9.688. n=60 neurons from four replicates. D3) Cytochalasin-B was added at DIV1 and S value measured at DIV3 (violin plot). 2-way ANOVA test, with post-hoc Tukey HSD. Row factor (Bl-/+): p<0.0001, F=29.03. Columns factor (Ctrl/Stretch): p<0.0001, F=36.07. n=60 neurons from four replicates.

However, it is well known that nuclear shape is also regulated by the perinuclear actin cap,[38] a shell-like structure that is connected to the INM through the LINC complex.[39] The actin cap transmits mechanical forces to chromatin, through a chain of interactions (actin cap / non-muscle myosin IIA (NMIIA) / emerin / lamin A/C).[40] We treated neurons with Blebbistatin (Bl, 0.4 µg/ml) to perturb the contractility of the actin filament bundles located in the cap, and we measured the *S* parameter. This dose did not interfere with *S* parameter of control neurons (p=1 for Bl+ vs Bl-in control nuclei) but it blocked the morphological changes induced by the stretching (p=0.92 for Bl+, ctrl vs stretch, Figure 3D2), suggesting that the actin cap is involved in force transmission from the cytoskeleton to the nucleoskeleton. To provide another line of evidence, we treated neurons with cytochalasin B (CB, 0.05 µg/ml), a drug that decreases the rate of actin polymerization[41]. The treatment *per se* does not alter the *S* parameter (p=0.48 for CB-vs CB+ in control nuclei) but nano-pulling loses its ability to remodel nuclear shape (p=0.25 for CB+, ctrl vs stretch, Figure 3D3).

### 2.3 Force transmission induces chromatin remodelling

In the preceding sections, we demonstrated that nano-pulling induces cytoskeletal remodeling and alterations in nuclear morphology. The observed correlation between these two events suggests a causal relationship, as all pharmacological treatments that modulate force generation at the level of microtubules and actin effectively abolish the nano-pulling effect. To further investigate force transmission from the cytoskeleton to the nucleus, we sought to determine whether the nuclei of stretched neurons experience greater tension compared to those of control neurons. To investigate this, we utilized a FRET (Förster resonance energy transfer) tension sensor developed by Conway and colleagues [42,43]. The sensor consists of an actin-binding domain and a SUN-binding domain of nesprin, which connect the cytoskeleton to the inner nuclear membrane, respectively. These two domains are separated by a FRET-based tension sensor, comprising two fluorescent proteins (mTFP1 and Venus) linked by a 40 amino acid elastic linker (Figure 4A1). When the NESPRIN sensor is under tension, the elastic linker is stretched, leading to a reduction in the FRET signal. We measured the FRET signal in neurons expressing the sensor by analysing both the medial (equatorial) and basal sections. As expected, we observed a decrease in FRET signal in stretched neurons compared to control neurons in the medial plane (E=0.37±0.01 in control and E=0.32±0.02 in stretched neurons, Figure 4A3). Since previous studies suggested that the basal plane show the greatest difference in signal,[42] we repeated the measurements in this plane. As anticipated, we found a more pronounced difference between the two groups (E=0.47±0.02 in control and 0.31±0.02 in stretched neurons, Figure 4A4).

**Figure 4.**
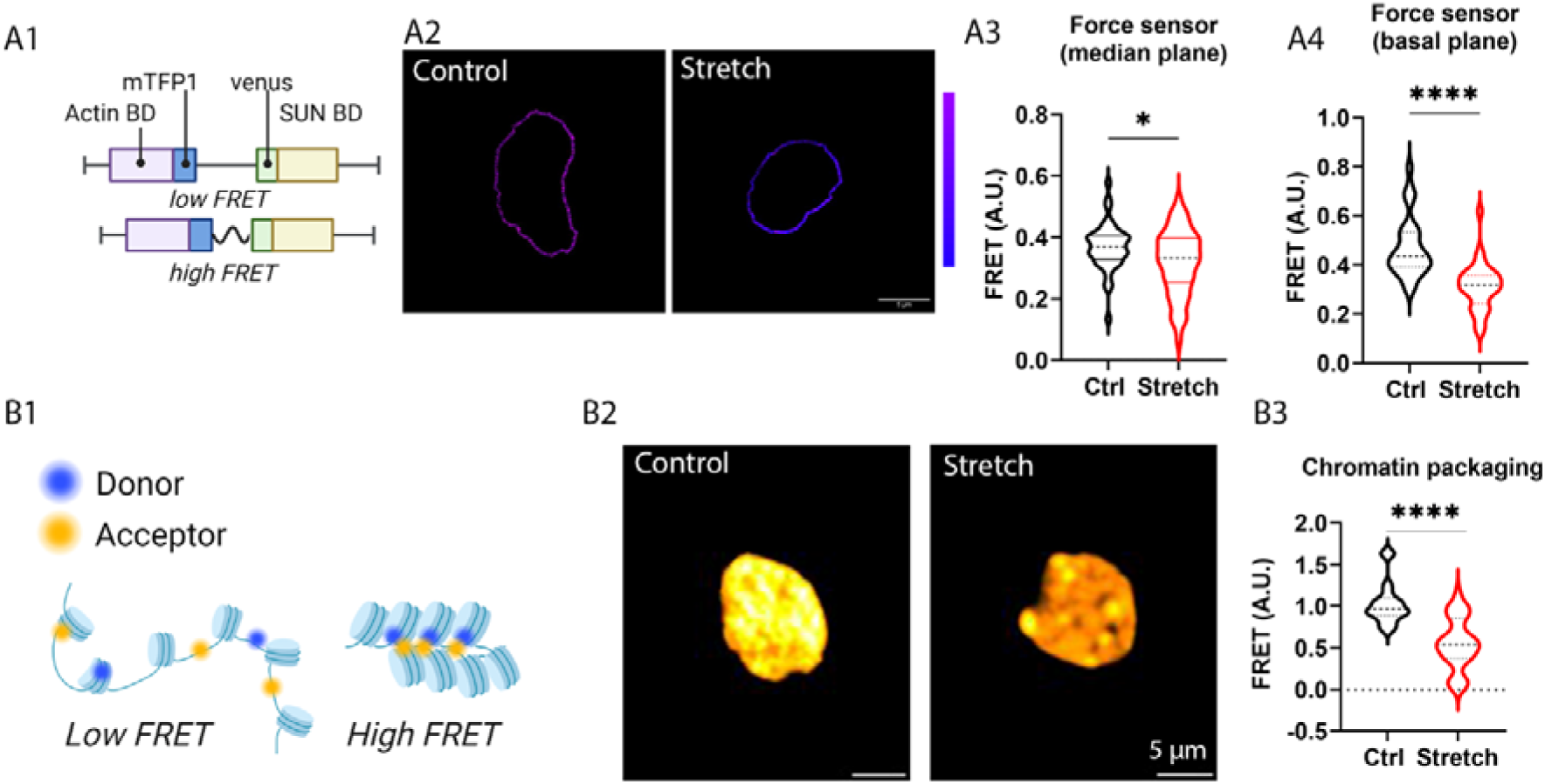
FRET sensors. A1) scheme the FRET tension sensor. A2) FRET signal from selected images, *scale bar* 5 µm. A3) FRET signal (median plane) data distribution (violin plot). Unpaired t-test test, two-tailed. P=0.01, t=2,591, df=86, n=45 neurons from four replicates. A4) FRET signal (basal plane) data distribution (violin plot). Mann-Whitney test, two-tailed, p<0.0001, U=102, n=30 neurons from four replicates. B1) FRET signal emitted from the donor/acceptor coupling correlates with chromatin packaging. B2) Selected images. B3) FRET signal data distribution (violin plot). Mann Whitney test, two-tailed. p<0.0001, U=127, n=30 neurons from six replicates.

Next, considering that the transmission of mechanical forces from the cytoskeleton to the nucleoskeleton can influence gene transcription, we investigated potential changes in chromatin remodelling. For this purpose, we utilized a quantitative FLIM-FRET (fluorescence lifetime imaging microscopy coupled with FRET) assay able to report on chromatin compaction.[44] This assay leverages the detection of FRET between two fluorescent DNA intercalants (Hoechst and SYTO13) as a proxy of chromatin packaging (Figure 4B1). Using this approach, we identified changes in chromatin organization at the level of nucleosome arrays. Nano-pulling induced a statistically significant decrease in the FRET signal (Figure 4B3, p<0.0001), suggesting a increased abundance of euchromatin in stretched neurons.

### 2.4 Force-mediated nuclear remodelling modulates global gene expression

Here, we investigated whether force-induced nuclear rearrangements also have direct implications for transcriptional modulation. RNA-seq was performed on hippocampal neurons in the control and stretched conditions to profile the whole transcriptome (n=6). The principal component analysis (PCA) showed that the two groups have distinct transcriptional profiles (Figure S3). We found 2227 differentially expressed genes (padj ≤ 0.01, |log2FoldChange|>1.5), and gene ontology (GO) was carried out to investigate which biological processes were dysregulated between the two conditions (Figure 5A, Figure S1, Fig S2). Many GO categories are related to gene expression regulation at the epigenetic level (chromatin remodelling, GO:0006338, fold enrichment, FE=1.76; chromatin binding, GO:0003682, FE=1.68), at the transcriptional level (regulation of DNA binding, GO:0051101, FE=2.87; DNA-binding transcription factor binding, GO:0140297, FE=2.04; transcription regulator complex, GO:0005667, FE=1.65, nuclear protein-containing complex, GO:0140513, FE=1.65) and post-transcriptional level (transcription elongation by RNA polymerase II, GO:0006368, FE=4.2; mRNA processing, GO:0022613, FE=1.85; RNA binding, GO:0003723, FE=1.57). The highest upregulated Panther GO-Slim molecular functions were RNA polymerase II complex binding (8 genes, FE=5.44, p=2.01E-03), microtubule binding (21 genes, FE=2.31, p=1.24E-03), chromatin binding (24 genes, FE=0.98, p=7.71E-04). The highest upregulated Panther GO-Slim cellular component is related to the mediator complex (8 genes, FE=4.08, p=1.78E-03).

**Figure 5.**
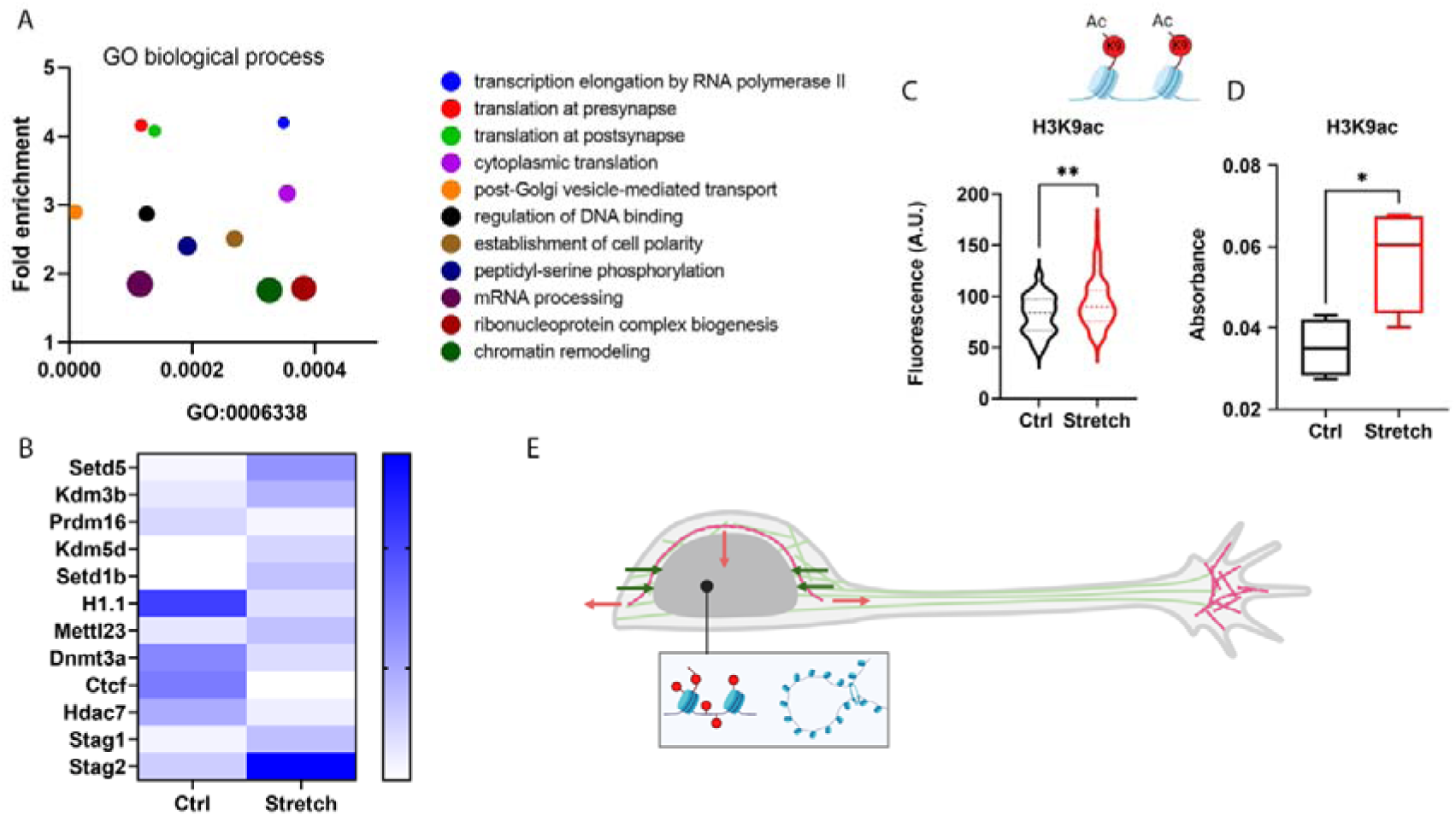
GO enrichment analysis (GOEA). A) GOEA via Panther (GO biological process), bubble plot (bubble size: number of genes). The bubble plot for GO cellular component and GO molecular functions are given in Figure S1 and Figure S2. Heat Plot map of DE (differentially expressed) genes belonging to GO:0006338 (chromatin remodelling). List of gene in chromatin organization (GO:0006325) related to histone/DNA methylation or histone acetylation or 3D chromatin architecture are also given in supplementary table S1. N=6 datasets per condition analysed. C) Global levels of H3H9 acetylation: estimated by IF analysis, Mann-Whitney test, two-tailed, n=90 nuclei from four replicates, p=0.0028. U value: 3011. D) Global levels of H3H9 acetylation: estimated by ELISA test, n=4 replicates, unpaired t test, two-tailed, p=0.0256, t=2.949, df=6. E) An integrated model of the cell remodelling induced by force. The green lines represent MTs. The red lines represent actin filaments. The red arrow indicates the contractile force generated by the actin network. The green arrows indicate the pulling force generated by MTs. The explanation is in the text.

In the gene list related to chromatin remodelling (GO:0006338), we identified many chromatin modifying enzymes (methylation, acetylation) and chromatin organizers (Figure5B, table S1). Interestingly, DEG (differential gene expression) analysis shows that eleven out of twelve of these factors are dysregulated in a way that favours transcriptional activation in stretched versus control neurons. We found an up-regulation of Setd1B, a lysine methyltransferase of H3K4,[45] a well-known epigenetic mark of active transcription enriched at the promoters / enhancers of highly expressed genes. We also found an up-regulation of the histone-arginine methyltransferase Mettl23, which catalyzes the demethylation of H3R17,[46] a modification that positively modulate transcription initiation.[47] We also found an up-regulation of the histone-lysine N-methyltransferase SETD5, a writer of H3K36me3,[48] an epigenetic mark enriched over coding regions of highly expressed genes. Conversely, H3K9me is a mark of transcriptional repression[49] that is recognized by the heterochromatin protein 1 (HP1). In this regard, we found an up-regulation of the lysine-specific demethylase Kdm3b that is an eraser of H3K9me,[50] and a down-regulation of the histone-lysine N-methyltransferase Prdm16, a writer of H3K9me.[51] We also found a downregulation of H1.1, a high mobile variant of histone H1 that promote the formation of higher order chromatin structures in vivo.[52] Extensive chromatin remodeling is also evidenced by the strong down-regulation of the insulator CCCTC-binding factor (CTCF), which plays a pivotal role in organizing chromatin loops and the 3D genome by stabilizing long-range interactions through multimerization.[53] The other major player in the formation of DNA loops is cohesion, a ring-shaped complex composed of a core trimer and one STAG subunits (STAG1 or STAG2, both up-regulated in our analysis). These cohesion complexes stabilize the promoter/enhancer interactions, being STAG1 more involved in long-range interactions and STAG2 in mid-range interactions.[54] We found down-regulation of other enzymes involved in transcriptional silencing, such as the histone deacetylase Hdac7 [55] or the DNA cytosine-5 methyltransferase Dnmt3a, an enzyme responsible for de novo DNA methylation that repress transcription by the establishment of a methylation patterns and recruiting deacetylases.[56]

To confirm that the nano-pulling positively modulates gene expression at the epigenetic level, we quantified the global acetylation levels of histone H3 at lysine 9 (H3K9Ac), a mark associated with active gene transcription. First, global levels of H3K9Ac have been quantified via immunofluorescence (IF), showing increased levels (estimated as mean fluorescence) in stretched neurons (Figure 5C). Second, we detected global levels of H3K9Ac by using an ELISA kit (Figure 5D). We observed a similar trend, but the increase was more pronounced (1.6-fold increase in the stretched versus control group), probably due to the higher sensitivity of this detection method.

## 3 CONCLUSION

Nuclei of adherent cells tend to have a flattened disk-like shape and smooth nuclear surfaces with no nuclear grooves. This is mainly due to the force generated by the actin cap that forms a curved shell tightly surrounding the apical surface of the nucleus, thereby controlling nuclear shape.[57] The actin cap is also physically connected to actin fibers and the perinuclear microtubules. Microtubules are force-bearing elements.[58] It is also well known that MTs can generate compressive forces onto the nucleus,[33] due to plus-end polymerization,[59] the relative sliding between MTs and the F-actin cortex (powered by dynein)[60] or the relative sliding between two MTs (powered by kinesin-1).[61] Nuclear shape is likely determined by the balance between the forces generated by the actin and microtubule cytoskeletons, respectively (see the schematic in Figure 5E). Perturbation of these forces could result in changes in nuclear shape.

In recent years, mechanical force has been recognized as an external factor that can stabilize MTs.[25–27,31,62,63] It has been proposed that MTs possess inherent mechanosensitivity, with computational models suggesting that tensile forces, even as small as a few piconewtons, can promote MT assembly. These forces are believed to facilitate the formation of lateral bonds between GTP-tubulin dimers, thereby aiding the closure of the protofilament wall,[64] and experimental data related to the modulation of MT dynamics in response to applied forces seem not to contradict this model.[65,66] Mechanical forces can also influence MT stability indirectly by affecting microtubule-associated proteins (MAPs), by modulating MT/MAP associations[67,68] or MAP activity[69] or MAP relocation.[70]

Recently, we demonstrated that the chronic application of a pulling force on the scale of tens of pico-Newtons induces the stabilization of MTs in developing axons.[27,31] This force-induced stabilization appears to be a general mechanism, as it was also observed in the neural processes of neural progenitor cells undergoing neural differentiation.[26] The data here presented confirm that stretched neurons have longer axons with more stable axonal MTs (Figure 1C), supporting previously published findings.[27] Here, the analysis was also extended to the soma, suggesting that force-induced MT stabilization is ubiquitous within the neuronal cytoplasm (Figure 1D).

However, while analysing the MT network at the somatic level, specifically in the perinuclear region, we uncovered evidence not previously reported. The orientation of perinuclear MTs changes in stretched neurons, with most of them aligning perpendicularly to the nuclear envelope (Figure 1A1-3, B1-2). This extended remodelling in response to applied force, never before described in neurons, closely resembles the changes observed in plant cells that reorient themselves in response to mechanical stress.[62,71–73]

In stretched neurons, we observed that perinuclear MTs oriented perpendicularly to the nuclear envelope indent the outer nuclear membrane (Figure 2A). The accumulation of nuclear grooves results in a statistically significant increase in the nuclear perimeter observed in stretched neurons (Figure 2B3). The more jagged contours cannot be attributed to changes in nuclear lamin B1(Figure 3C). We speculate that MTs generate a force that locally stretches the nuclear envelope, likely causing nuclear grooves. To investigate whether the nuclear envelope is under lateral compression in stretched neurons, we leveraged a FRET tension sensor consisting of two domains of nesprin, which connect the cytoskeleton to the inner nuclear membrane (INM). Our data show a decrease in the FRET sensor signal, indicating that nesprin, and consequently the nuclear envelope, is under stretching in the horizontal plane (Figure 4A). Considering that the MT network is physically connected to the actin cup, it is reasonable to infer that the MT-transmitted pushing force not only contributes to the formation of nuclear grooves but also partially counteracts the pulling force generated by the actin cup. This is suggested by the evidence that the nuclei of stretched neurons drastically change their shape (Figure 3B), becoming less flat, as documented by the increase in height along the Z plane, the decrease in the cross-sectional area of the medial plane, and the increase in sphericity (Figure 3C). This morphological change may also contribute to the appearance of grooves on the nuclear surface. We observed that treatment with nocodazole, a pharmacological agent that impairs microtubule (MT) stabilization, prevents force-induced changes in nuclear morphology (Figure 3D1). Interestingly, treatment with blebbistatin (an inhibitor of actin-myosin interaction) or cytochalasin B (an inhibitor of actin filament formation) also prevents nuclear shape changes in response to force (Figure 3D2-3). One possible explanation for these observations is that when the MT polymerization is inhibited or MT/actin coupling and actin-mediated force transmission are disrupted, MTs can no longer function as load-bearing elements to counteract the pulling force generated by the actin cup. Our findings related to the ability of MTs to change nuclear shape are also consistent with a previous observation that, after pharmacological disruption of microtubule, the polymerization of the newly formed microtubules reshapes the nuclear envelope and the chromatin fibres inside the interphase nucleus.[60]

Forces propagated from the cytoskeleton to the nesprins can be further transmitted to the nuclear lamina, which serves as the primary load-bearing element within the nucleus.[74] Studies on purified nuclear lamins *in vitro* have demonstrated that these proteins can withstand substantial mechanical forces, deform under stress, and exhibit strain-stiffening properties.[75–77] Inside the nucleus, the LINC complex connects to the nuclear lamina, which anchors large chromatin loops. In addition to their localization at the nuclear envelope, A-type lamins are distributed throughout the nucleoplasm, where they assemble and interact with chromatin.[76] Lamin proteins bind to core histones through their C-terminal tails, facilitating force transmission between the lamins and chromatin.[78,79] It has been reported that mechanical forces can be conveyed through the nuclear lamina to the chromatin,[80] influencing both chromatin organization[81] and gene expression.[82,83] To test this hypothesis in our model, we used a FRET sensor to evaluate chromatin condensation. We observed a decrease in global chromatin packaging (Figure 4B), which is associated with the activation of gene transcription. These findings are consistent with previous studies showing that intracellular force transmission to the nucleus triggers a decrease in chromatin compaction, chromatin redistribution, loss of repressive mark and activation of transcriptional activity.[34,84,85] To confirm this data, we evaluated global levels of H3K9ac, an epigenetic mark associated with active transcription. Using two independent assays - immunofluorescence (IF) and ELISA detection - we found a global increase in the levels of this epigenetic mark in stretched neurons (Figure 5C, D). To further validate this, we performed RNA sequencing (RNA-seq) to profile the global transcriptome in two conditions (stretched versus control neurons). Subsequent analysis (GO, GOEA) revealed that, in terms of fold enrichment score, the most relevant cellular component is the activation of local translation, particularly at the synapses. This is consistent with our previous reports where we demonstrated that force induces activation of local translation.[27,31] In terms of gene count, the most relevant cellular processes are those related to chromatin remodelling (GO:0006325). Figure 5B lists the most differentially expressed genes, most of which are readers and writers of specific epigenetic marks (H3K4me, H3K36me3, H3K9me, 5mC), histone variants (H1.1), insulators (CTCF), and proteins involved in organizing chromatin loops and 3D topologically associated domains (STAG1 or STAG2). Of note, most of them (11 out of 12) correlate with transcription activation (table S1).

In conclusion, we propose a model (Figure 5E) in which force induces the stabilization of microtubules. In the soma, the stabilization of MTs induces a reorganization and reorientation of the MT network, similar to what has already been described and well-characterized in plant cells.[71] Biophysically, the pulling force generated by the perinuclear MTs (green arrows in the scheme) relaxes the contractile force produced by the actin cap (red arrows in the scheme), which tends to flatten the nucleus. As a result, the nucleus becomes more spherical, and surface grooves appear. At the level of gene expression regulation, the force generated by MTs propagates to the nuclear envelope. Since chromatin is mechanically tethered to the nuclear envelope through lamins, this force propagation induces a global chromatin remodelling, reducing chromatin packaging and resulting in global transcriptional activation. This model is consistent with *in vivo* observations that fluctuations of the nuclear envelope, induced by MTs, lead to a broad ‘stirring’ of the genome, enhancing chromatin accessibility and stimulating gene expression.[86]

## 4 Experimental Section

### 4.1 Data availability

Authors declare to adhere to OS (open science) principles. Raw data, metadata, processed data associated to this publication are available in Zenodo repository (10.5281/zenodo.13961112). RNA-seq data have been deposited at GEO at GEO: accession number GSE197808.

### 4.2 Animals

All animal procedures were performed in compliance with protocols approved by Italian Ministry of Public Health and of the local Ethical Committee of University of Pisa (protocol number 39E1C.N.5Q7), in conformity with the Directive 2010/63/EU. For this work, C57BL/6J mice (post-natal day, P1) were used. Animals were kept in a regulated environment (23 ± 1°C, 50 ± 5% humidity) with a 12/12 h light/dark cycle and food and water *ad libitum*.

### 4.3 Cell culture

Hippocampal neurons were isolated from P1 mice (both sexes). Animals were sacrificed by cervical dislocation and both hippocampi were harvested in a solution of D-glucose 6.5 mg/ml in DPBS (Gibco, Thermo Fisher Scientific, Waltham, Massachusetts, US, #14190-094). Then, chemical and mechanical dissociation was performed to isolate the primary neurons, following a protocol already published [87]. Neurons were seeded in high-glucose DMEM (Gibco, Thermo Fisher Scientific, Waltham, Massachusetts, US, #21063-029) with 10% fetal bovine serum (FBS, Gibco; Thermo Fisher Scientific, Waltham, Massachusetts, US, #10270-106), 100 IU/ml penicillin, 100 μg/ml streptomycin (Gibco, Thermo Fisher Scientific, Waltham, Massachusetts, US, #15140-122) and 2 mM Glutamax (Gibco, Thermo Fisher Scientific, Waltham, Massachusetts, US, #35050-038) on surfaces pre-coated with 100 μg/ml poly-L-lysine (PLL, Sigma-Aldrich, Burlington, Massachusetts, US, #P4707). After four hours, the medium was replaced by a cell culture medium consisting of Neurobasal-A medium (Gibco, Thermo Fisher Scientific, Waltham, Massachusetts, US, #12348-017) modified with B27 (Gibco, Thermo Fisher Scientific, Waltham, Massachusetts, US, #17504-044), 2 mM Glutamax, 50 IU/ml penicillin, 50 μg/ml streptomycin. To magnetically actuate neurons, superparamagnetic iron oxide nanoparticles (SPIONs) were added to the cell culture medium with a final concentration of 5 µg/ml. Cultures were maintained at 37°C in a saturated humidity atmosphere of 95% air and 5% CO_2_.

### 4.4 Drug treatment

Neurons were cultured in canonical cell culture medium from DIV0 to DIV1. At DIV1, the medium was supplemented with Nocodazole (Sigma-Aldrich, Burlington, Massachusetts, US, #SML1665; 1.8 ng/ml), Cytochalasin-B (Sigma-Aldrich, Burlington, Massachusetts, US, #C6762; 0.05 µg/ml) or Blebbistatin (Sigma-Aldrich, Burlington, Massachusetts, US, #B0560; 0.4 µg/ml). The drug concentrations used in this work have been already validated on hippocampal neurons for not inducing morphological changes in axon growth.[25]

### 4.5 Plasmid Amplification and Primary Cell Transfection

The pcDNA nesprin TS plasmid was purchased by Addgene (plasmid #68127; http://n2t.net/addgene:68127; RRID). Bacteria obtained from Addgene were streaked onto agar plates containing the appropriate antibiotic. Plasmid DNA was subsequently amplified and isolated from 100 mL of cultured bacteria using the Macherey-Nagel NucleoBond Xtra Midi kit (Macherey-Nagel, #740412).

Transfection was performed using Lipofectamine 2000 (Invitrogen, #11668), with modifications aimed at reducing cellular stress. Briefly, 4 × 105 primary hippocampal neurons were seeded onto a 35-mm Quad ibidi μ-Dish (Ibidi GmbH, #80416), pre-coated with PLL and laminin (0.002 mg/mL; Sigma, #P2020), using the appropriate growth medium. After allowing the cells to attach for two hours, 0.5 μg of total plasmid DNA per well was used for transfection, according to the manufacturer’s instructions. Following a two-hour incubation at 37°C in the CO2 incubator, the transfection mixture was removed, and the cells were returned to culture medium.

### 4.6 Nano-pulling

To generate magnetic nano-pulling, 100 nm SPIONs (Fluid-MAG-ARA, Chemicell, Germany; #4115) were used. Specifically, SPIONs are particles characterized by an outer layer of glucoronic acid, to prevent aggregation, surrounding a core of iron oxide (approximately 75 ± 10 nm in diameter). As stated from the supplier, they present a saturation magnetization of 59 A·m^2^/kg. To provide the external magnetic actuation, A Halbach-like cylinder magnetic applicator was used. Such an applicator generates constant magnetic field gradient of 46.5 T/m in the radial centrifugal direction as previously demonstrated.[88,89] At DIV0, hippocampal neurons were seeded with a density of 160 cells/mm^2^ - unless stated otherwise - and were treated with 5 µg/ml SPIONs four hours from the seeding (DIV0.17). The day after (DIV1), SPION-modified samples were split into controls (null magnetic field) and stimulated (stretch group). Magnetic stimulation was applied on a time-scale ranging from 48 hours to 120 hours.

### 4.7 Micro-fluidic system

Gene expression studies have benefited from the use of a system of microfluidic chambers, suitable for separating axonal and somato-dendritic compartments. To do this, we used XONA microfluidic devices (XONA, Research Triangle Park, North Caroline, US, #RD150) previously mounted on glass coverslips (22 mm in diameter). To promote the separation between neuronal compartments – and thus pushing axons to cross the micro-channels – we applied a different coating. Briefly, in the somato-dendritic compartment we applied the canonical coating, while in the axonal compartment 500 μg/ml PLL and 100 μg/ml laminin were provided.

### 4.8 RNA extraction and quantification

RNAs from somato-dendritic compartments, in control and stretch groups, were extracted following a protocol already published.[90] Briefly, RNA extraction was performed with nucleoSpin RNA PLUS XS Kit (Machery-Nagel, Düren, Germany, #740990.50). Samples were collected in lysis solution and incubated at 4°C for subsequent steps, following manufacturer’s protocol. RNA was eluted in RNAse-free water and quantified with the Quant-iT RiboGreen RNA Kit (Thermo Fisher Scientific, Waltham, Massachusetts, US, #R11490).

### 4.9 RNA sequencing

RNA sequencing was performed on stretch and control groups after 120 hours of magnetic nano-pulling (DIV6). Six biological replicates, for a total of 12 extracts, were analysed. Quality check (QC) was performed by calculating the ribosomal content (RNA integrity number). For sequencing, the platform Illumina NovaSeq6000 sequencer was used (GENOME Scan, The Netherlands). The RNA library was prepared using the polyA selection library and sequencing mode PE (paired end), read length 150 bp, ∼9 Gb and 30 million XP reads per sample. Default genomes and mouse annotation were Ensembl GRCm39. BAM data, gene count and differential expression (DE) analysis were performed on Galaxy platform (https://usegalaxy.eu/) with BWA-MEM, featureCounts ad DEseq2 tools, respectively. The gene ontology GO and GO enrichment analysis GOEA was performed by PANTHER Overrepresentation Test (DOI: 10.5281/zenodo.7942786) the analysed list was mus musculus (all genes in database), FISHER test, FDR correction, GO biological process.

### 4.10 ELISA

At DIV3, mouse hippocampal neurons were collected and leveraged to quantify H3K9 acetylation levels by ELISA assay. To do this, PathScan® Acetylated Histone H3 Sandwich Elisa Kit (Cell Signaling Technology, Massachusetts, US, #7232C) was used. Briefly, hippocampal neurons were seeded at DIV0 on 30 mm petri dishes and cultured for up to 48 hours. Then, cells were homogenised in lysis buffer (Cell Signaling Technology, Massachusetts, US, #9803) to which phenylmethylsulphonyl fluoride (PMSF) was added. The samples were incubated in this solution for 5 minutes and then sonicated. By centrifugation (14000 prm for 10 minutes) the supernatant was separated and used for subsequent steps. Protein concentration was quantified by Pierce^tm^ BCA Protein Assay Kit (Thermo Fisher Scientific, Waltham, Massachusetts, US, #23225). Diluent X (Cell Signaling Technology, Massachusetts, US, #11083) was added to the supernatant to achieve a final volume of 100 ul. Subsequent incubations with primary and secondary antibodies were carried out following manufacturer’s protocol. Absorbance was measured by a plate reader set at 450 nm and H3K9 ac was quantified according to the standard curve.

### 4.11 Immunostaining

Hippocampal neurons were fixed in a solution of 2% PFA, 7.5% sucrose for 20 minutes at RT. Fixed samples, were first permeabilized in 0.5% Triton X-100 for 10 min at RT and then blocked in 5% serum, 0.3% Triton X-100 in DPBS for 1 hour at RT. Samples were incubated ON at 4°C with primary antibodies (TUBB3, Sigma-Aldrich, Burlington, Massachusetts, US, #T8578, 1:500; acetylated tubulin, Sigma-Aldrich, Massachusetts, US, #T7451, 1:400; tyrosinated tubulin, Abcam, # ab6160, 1:400; acetylated H3K9, Cell Signaling Technology, Massachusetts, US, # C5B11, 1:200; αLaminB1,Proteintech, #66095-1-Ig, 1:500). The day after, samples were incubated for 1 hour at RT with appropriate secondary antibodies (Thermo Fisher Scientific, Waltham, Massachusetts, US, #O6380, #A32731, #A11011, #A32728, #R6393, #A21449, 1:500) and Hoechst 33342 (Thermo Fisher Scientific, Waltham, Massachusetts, US, #H3570, 1:1000). Images were acquired with a laser scanning confocal microscope (Nikon A1, Eclipse Ti) and super-resolution microscopy (Airyscan 2, Zeiss) or single module detection (SMD) confocal microscope (Leica SP5), by a 60× objective oil immersion. For acquisition, a 405 nm laser (425–475 emission filter) or a 488 nm laser (500– 550 emission filter) or a 561 laser (570–620 emission filter) or a 640 nm laser (663–738 emission filter) was used. Gain and exposure time varied for the specific experiments and the related information are available in the images that can be downloaded in the original format .nd2 from ZENODO repository (10.5281/zenodo.13961112).

### 4.12 Image analysis

For fluorescence quantification, ImageJ software was used.[91] Mean fluorescence *(f^-^)* was evaluated following the formula reported here:[27]

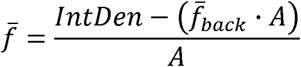

Where A is the area of the ROI (region of interest), IntDen is the integrated density (defined as the sum of all the pixel intensities in the ROI), and *(f^-^_back_)* is the mean fluorescence of the background readings (at least 3).

To determine the stability of MTs under the two conditions, we evaluated the ratio between acetylated and tyrosinated α-tubulin, intended as the ratio between the respective *(f^-^)*. To assess the area of the nucleus cross-section (S), we built the ROI of the nucleus by using Hoechst 33342 and then we calculated the extension in ImageJ.

For nuclear morphology, after lamin B1 immunostaining, images were acquired at 60X magnification, with an additional 4X digital magnification. Z-stacks were taken with a step size of 0.195 µm. For 2D reconstruction, “Z project” command was used. For the perinuclear ring, the ROI of the nucleus was selected as outer ROI and the “enlarge” command (−0.5µm) was used to selected the inner ROI. For 3D reconstruction, the ImageJ 3D Suite plugin was used.[35] The ROI of the nucleus was selected, and the Gaussian Blur 3D filter was applied, using standard settings to smooth intensity. Next, nuclei segmentation was performed using the Otsu threshold, a command within the 3D Suite plugin. At this stage, any holes left unprocessed by the segmentation were manually filled. Finally, the following tools from the 3D Suite plugin were applied: Volume, Surface, 3D Feret, 3D Compactness, and 3D Ellipsoid.

### 4.13 Fluorescence Resonance Energy Transfer (FRET) and Fluorescence Lifetime Imaging Microscopy and FRET (FLIM-FRET)

To evaluate the state of chromatin compaction, FLIM-FRET experiments were performed. At DIV3, hippocampal were fixed and stained for TUBB3 antibody, following the protocol described above. The need to mark samples with β-tubulin III stems from the necessity to distinguish neuronal cells in a culture that is not completely pure. FRET technique is based on energy transfer, i.e., a distance-dependent physical process by which energy is transferred from an excited molecular fluorophore (donor) to another fluorophore (acceptor).[92,93] For this, Hoechst 33342 and SYTO13 (Thermo Fisher Scientific, Waltham, Massachusetts, US, #S7575) were used as donor and acceptor, respectively according to the procedure reported in.[44] In short, in each experiment PFA-fixed cells were first loaded for 15-30 min by Hoechst 33342 (2 mM). Lifetime of the donor in the nucleus was collected in 5-10 cells by a Leica SP5 SMD confocal microscope equipped with a PicoHarp 200 (Picoquant, Berlin) card. Then, SYTO13 (2-4 mM) was added to the cell buffer. After 15-30 min the lifetime of the donor in the nucleus was collected in 10-15 stretched cells. Acquisition parameters were set as follows: 405 nm pulsed excitation (40 MHz, Picoquant laser), 256×256 pixel with 1000 line/s scanning rate, 120 s total acquisition. Donor signal was acquired by a spectral detector collecting 430-500 nm photons. Donor cross-talk and acceptor (SYTO13) signal were acquired by a spectral detector collecting 540-600 nm photons. Acquisition of both channels was concomitant. Data elaboration was carried out as described in[44] by using SimFCS (Laboratory for Fluorescence Dynamics, University of California, Irvine) to generate the phasor plot and a custom-made FiJI plugin (available on request) to calculate FRET.

For the NESPRIN sensor, at DIV3, hippocampal were fixed and stained for αLaminB1, following the protocol described above. Image we acquired according to the procedure reported in.[42]

### 4.14 Sample preparation for ultrastructural characterization

Briefly DIV 3 primary hippocampal neurons were fixed with 1.5% Glutaraldehyde in sodium cacodylate buffer (0.1 M; pH 7.4), as monolayer, for 1 h at room temperature. Next, cells were manually scraped, collected and centrifuged for 15 min in the same fixative solution until a visible pellet was obtained, and kept in fresh fixative solution overnight at 4 °C. Cells were then post-fixed with reduced osmium tetroxide (1% OsO4 and 1% K_3_Fe(CN)_6_ in sodium cacodylate buffer (0.1 M; pH 7.4)). Samples were then rinsed, en bloc stained for 1 h, with pure X solution (Moscardini et al., 2020) as contrast agent. After the dehydration, with a growing series of ethanol concentration, cells were finally embedded in epoxy resin (Epon 812, Electron Microscopy Science, Hatfield, PA, USA) that was then baked for 48 h at 60 °C. For the ultrastructural analysis, 90-nm thin sections were cut using a UC7 (Leica Microsystems, Vienna, Austria) and collected on 300 mesh copper grids. The analysis was done using a Zeiss Libra 120 Plus transmission electron microscope operating at 120 kV and equipped with an in-column omega filter and 16-bit CCD camera 2k × 2k bottom mounted.

### 4.15 TEM image analysis

Multiple images (between 4 to 36) have been acquired at the magnification of 8000 X, and automatically stitched using Image SP Viewer software, to build up the whole nuclear compartment. These images have been used both for the analysis of axes length, nuclear perimeter, and area but also for the analysis of MTs’ orientation in the perinuclear region. The nuclear contour has been traced as a closed line by using NeuronJ, a Fiji plugin.[94] The corresponding ROI as has used to calculate the “convex hull” from the selection tool (the convex hull of a set of points X in Euclidean space is the smallest convex set containing X). The perimeter of the ROI corresponding to the nuclear contour (*P*_n_ and of the ROI corresponding to the convex hull *P*_ch_ were then calculated. The ratio α = *P_n_/P_ch_* was used as a mark of the nuclear grooves.

The nuclear envelope has been manually encircled and measures of Area and Perimeter have been mesured with imageJ software.

The orientation of MTs has been evaluated, in respect to the nuclear envelope (green line in Figure2A), as parallel or perpendicular (red line in Figure2A). MTs that are orthogonal to the cutting plane (red spot in Figure2A) are nor considered for the orientation. Several images (18 over 76 for each experimental point) did not show an evident orientation for MTs and so not included in the quantification.

Please provide precise details of all the procedures in the paper (behavioral task, generation of reagents, biological assays, modeling, etc.) such that it is clear how, when, where, and why procedures were performed. We encourage authors to provide information related to the experimental design as suggested by NIH and ARRIVE guidelines (e.g., information about replicates, randomization, blinding, sample size estimation, and the criteria for inclusion and exclusion of any data or subjects).

### 4.16 Statistical analysis

Data were plotted with GraphPad 7.0. Values are reported as the mean ± standard error of the mean (SEM). The normality of the distribution was tested using Kolmogorov-Smirnov normality test. For normally distributed data, we used the t test for unpaired data or ANOVA test followed post-hoc Tukey HSD. For non-normally distributed data, the Mann-Whitney test was carried out. Significance was set at p ≤ 0.05. Sample size estimation was estimated with G-power. Chi-square test was used for categorical data Figure legends include the statistical tests used, exact value of n, what n represents confidence intervals, and test values. Experiments were analysed in blind by two different operators.

## Supporting Information

Supporting Information is available from the Wiley Online Library.

## Acknowledgements

The study was partially supported by the Wings for Life Foundation (WFL-IT-16/17 and 20/21), the EC programme Horizon2020 (101007629-NESTOR-H2020-MSCA-RISE-2020), the Human Frontier Science Program (RGP0026/2021), and the European Union Next-GenerationEU - National Recovery and Resilience Plan (NRRP) – mission 4 component 2, investiment n. 1.4 – CUP N. B83C22003930001 (Tuscany Health Ecosystem – THE”, Spoke 8). This manuscript reflects only the authors’ views and opinions, neither the European Union nor the European Commission can be considered responsible for them. We would like to thank the Center for Instrument Sharing of the University of Pisa (https://cisup.unipi.it/). Received: ((will be filled in by the editorial staff)) Revised: ((will be filled in by the editorial staff)) Published online: ((will be filled in by the editorial staff))

---

Nano-pulling induces the stabilization, reorganization, and reorientation of the MT network. The nucleus of stretched cells experiences higher lateral tension, becomes more spherical, and develops surface grooves. At the level of gene expression regulation, changes in nuclear morphology induce global chromatin remodeling, reducing chromatin packaging and resulting in widespread transcriptional activation.

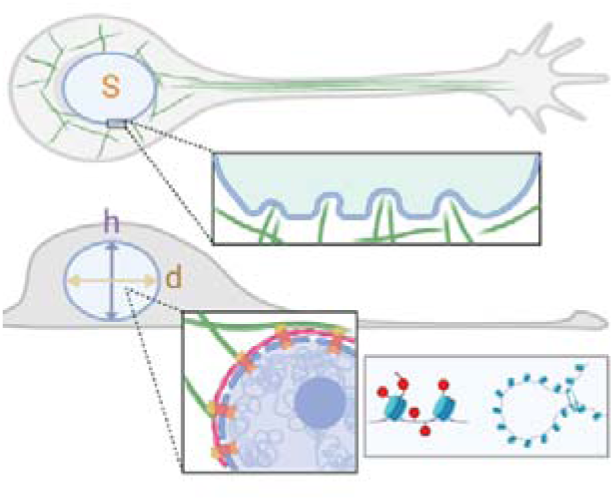

## Supporting Information

**Figures S1:**
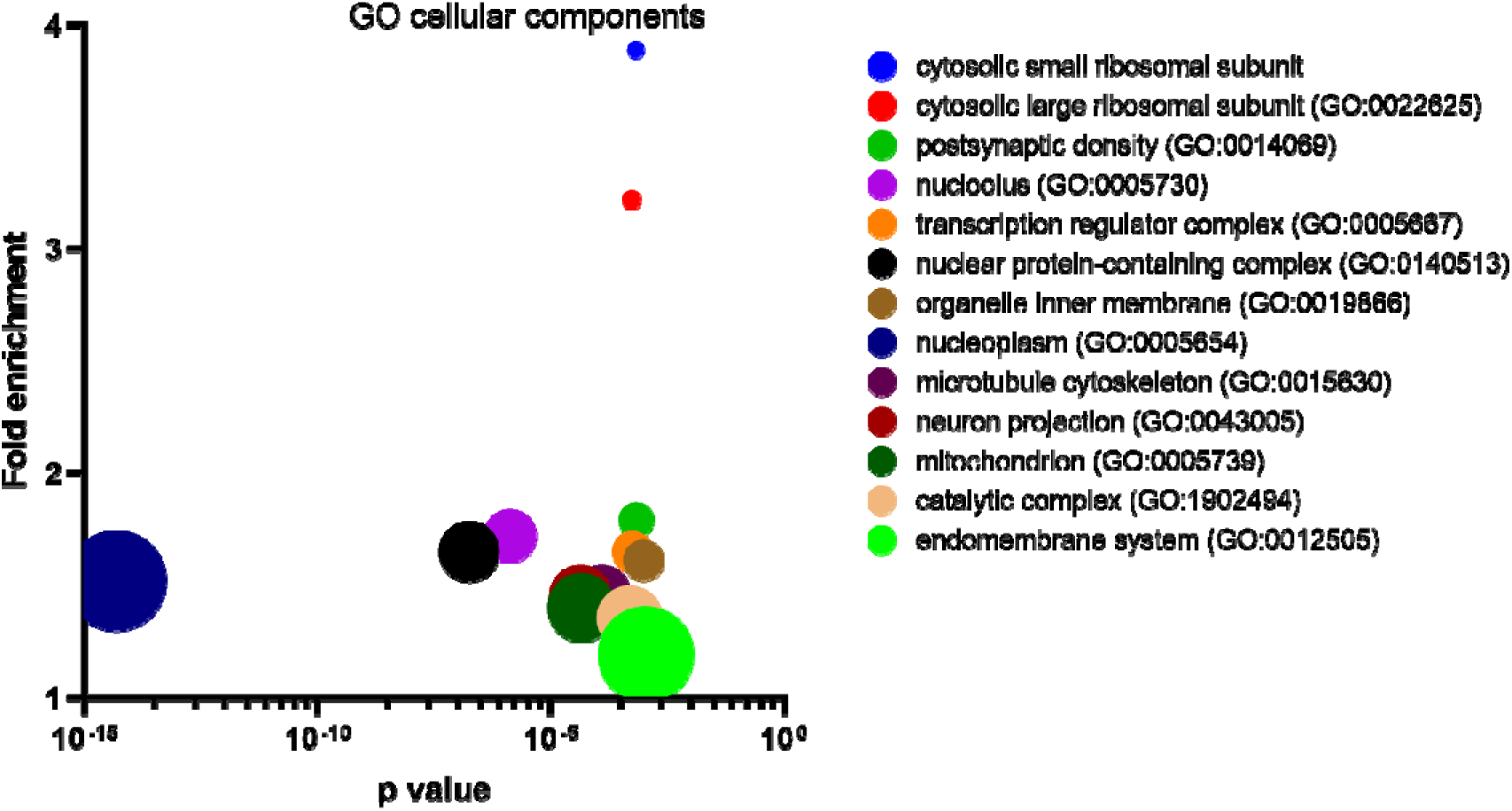
GOEA via Panther (GO cellular component), bubble plot (bubble size: number of genes).

**Figures S2:**
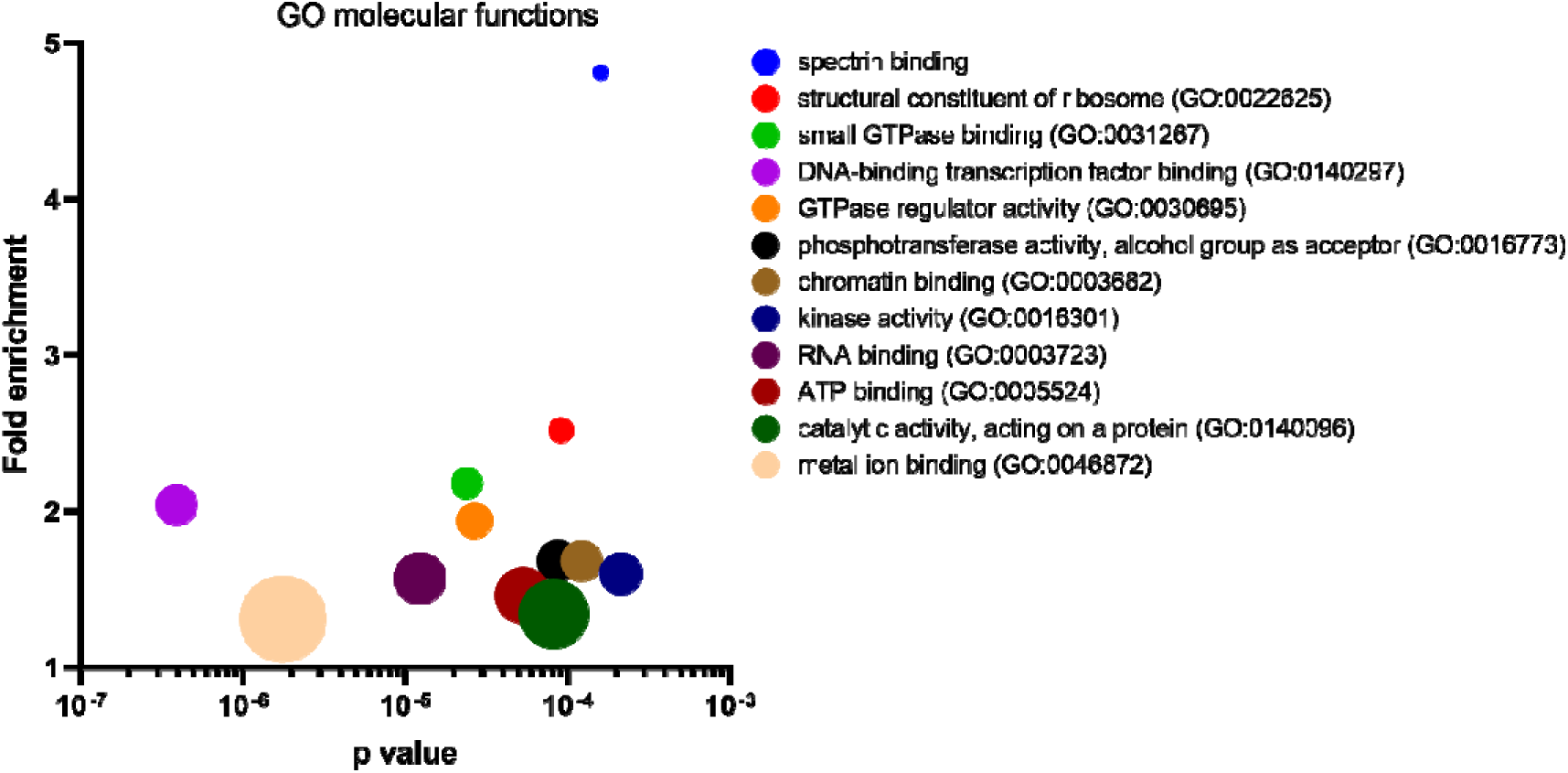
GOEA via Panther (GO molecular functions), bubble plot (bubble size: number of genes).

**Figures S3:**
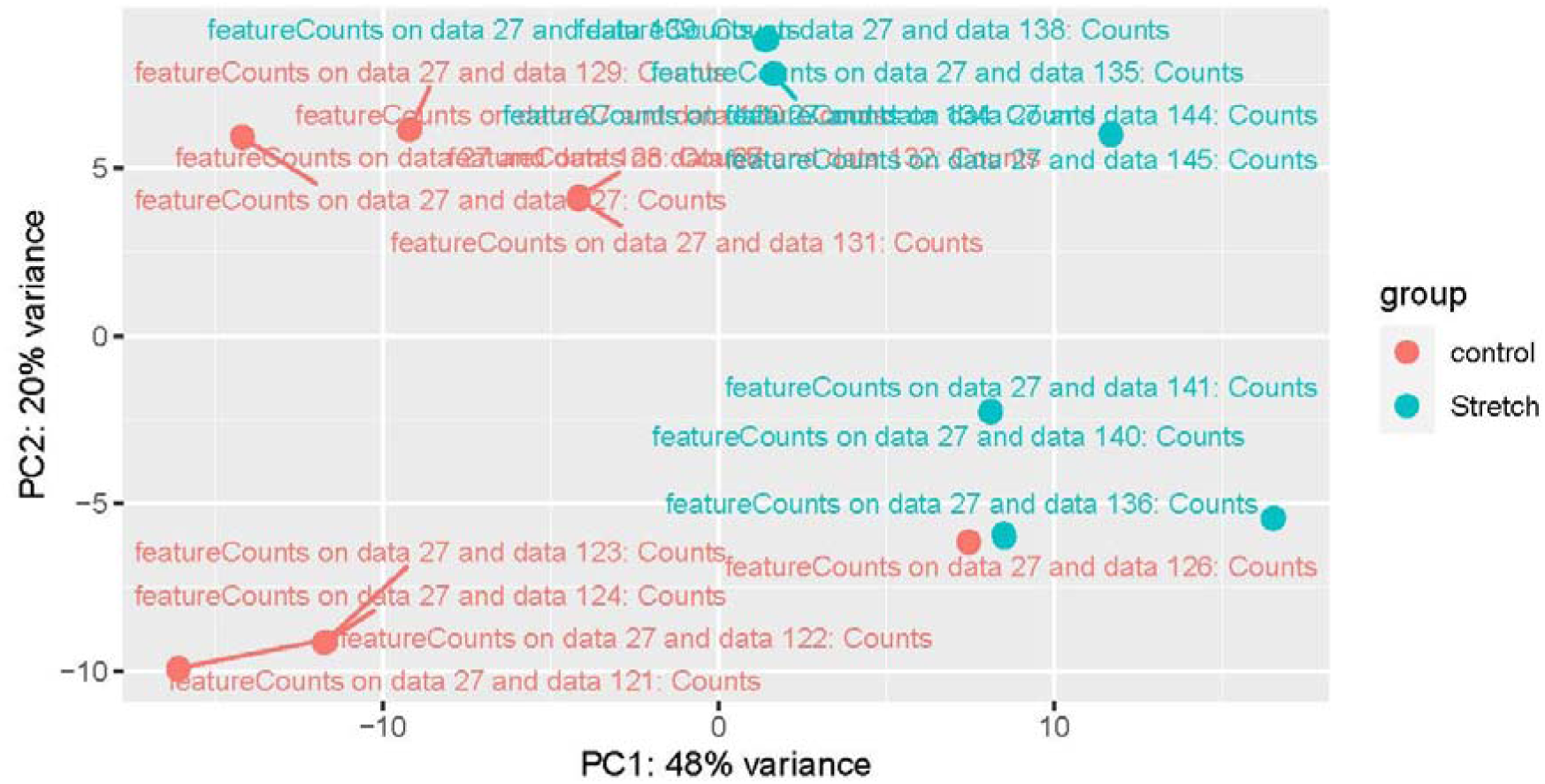
principal component analysis (PCA) from RNA-seq.

**Figures S4:**
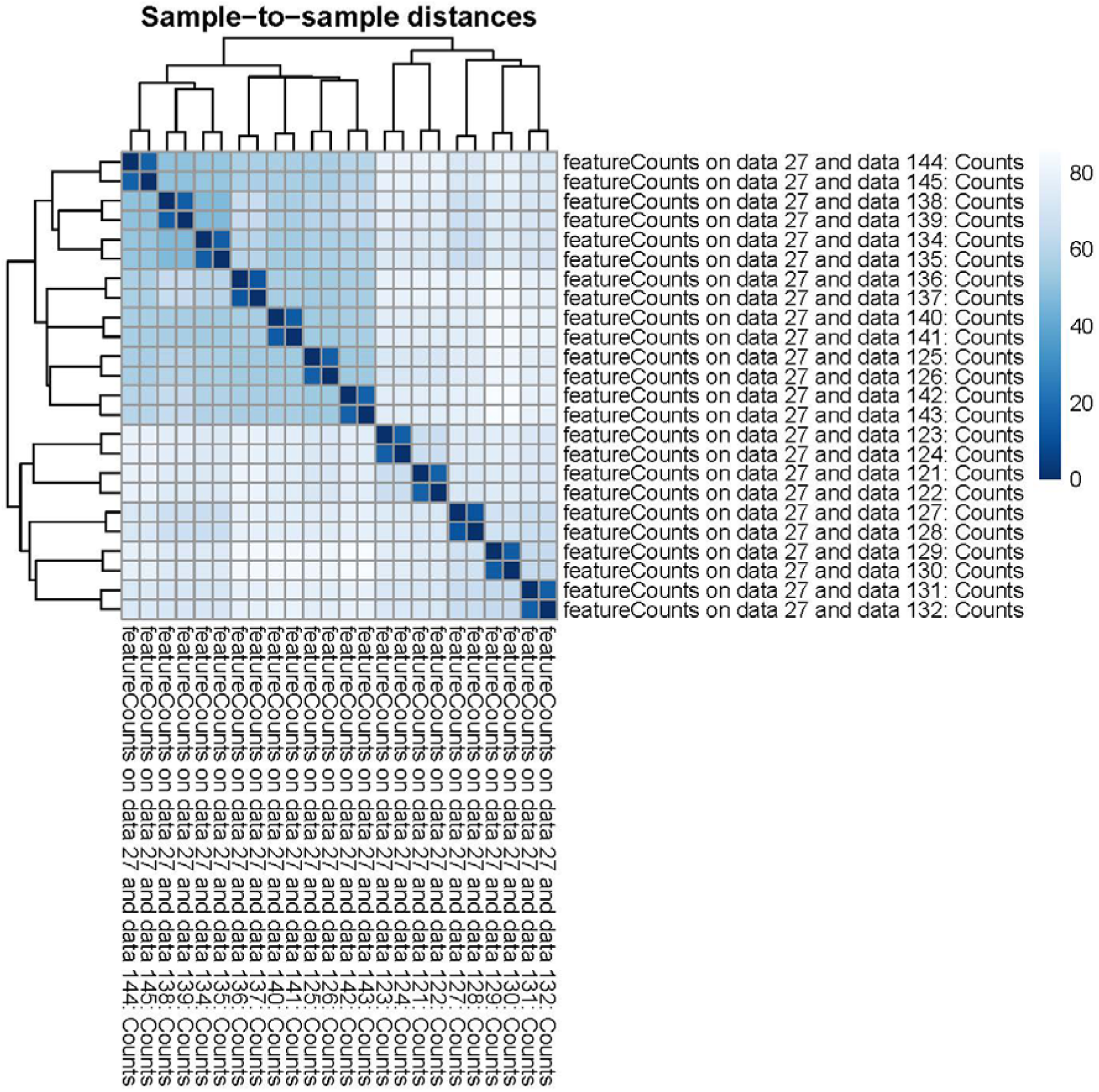
sample-to-sample distance from RNA-seq.

**Figures S5:**
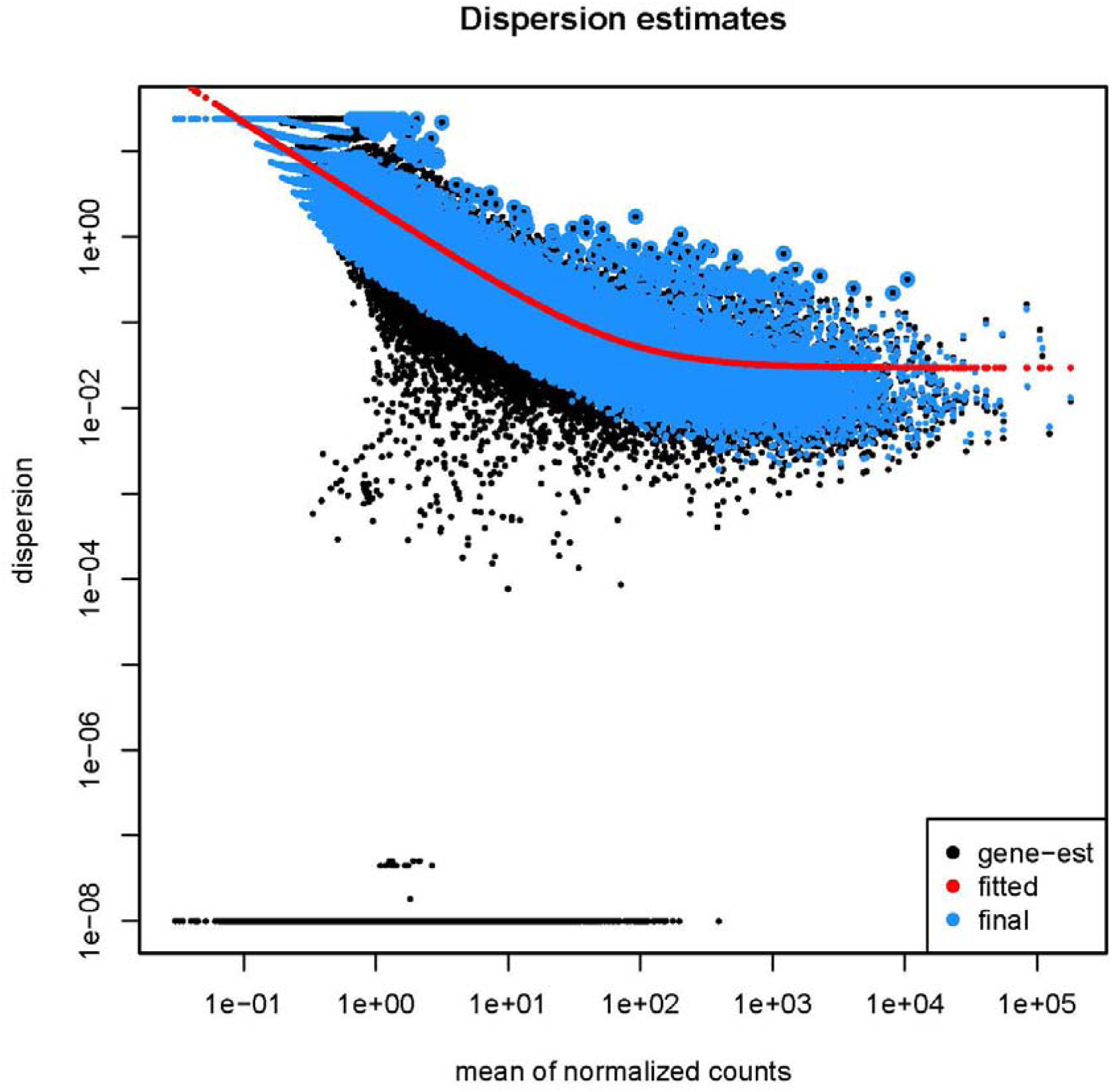
dispersion plot from RNA-seq.

---

**Table S1:**
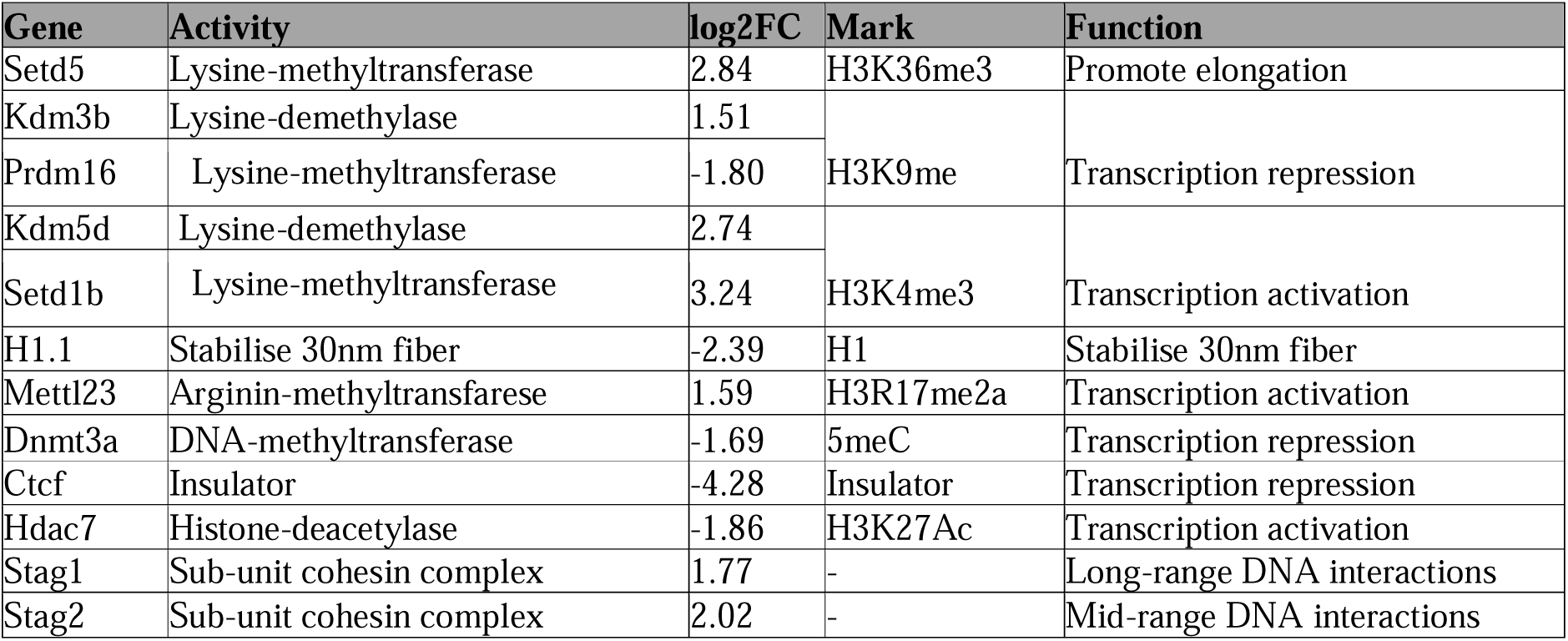
gene list (GO:0006325)

**Table S2.** Reagent list.

| Reagent/ Resource | Reference or Source | Identifier or Catalog Number |
| --- | --- | --- |
| **Experimental Models** |  |  |
| C57BL/6 mice, post-natal day (P1) | Charles River | 027C57BL/6 |
| **Antibodies** |  |  |
| TUBB3 | Sigma-Aldrich | #T8578 |
| Acetylated tubulin | Sigma-Aldrich | #T7451 |
| Tyrosinated tubulin | Abcam | # ab6160 |
| Acetylated H3K9 | Cell Signaling Technology | C5B11 |
| αLaminB1 | Proteintech | #66095-1-Ig |
| Goat anti-Chicken IgY (H+L) Secondary Antibody, Alexa Fluor™ 647 | Thermo Fisher Scientific | #A21449 |
| Goat anti-Mouse IgG (H+L) Secondary Antibody, Oregon Green 488 | Thermo Fisher Scientific | #O6380 |
| Goat anti-Rabbit IgG (H+L) Highly Cross-Adsorbed Secondary Antibody, Alexa Fluor™ Plus 488 | Thermo Fisher Scientific | #A32731 |
| Goat anti-Rabbit IgG (H+L) Cross-Adsorbed Secondary Antibody, Alexa Fluor™ 568 | Thermo Fisher Scientific | #A11011 |
| Goat anti-Mouse IgG (H+L) Highly Cross-Adsorbed Secondary Antibody, Alexa Fluor™ Plus 647 | Thermo Fisher Scientific | #A32728 |
| Goat anti-Mouse IgG (H+L) Cross-Adsorbed Secondary Antibody, Rhodamine Red™-X | Thermo Fisher Scientific | #R6393 |
| Hoechst 333 | Thermo Fisher Scientific | #H3570 |
| SYTO13 | Thermo Fisher Scientific | #S7575 |
| **Chemicals, reagents** |  |  |
| DPBS | Gibco, Thermo Fisher Scientific | #14190-094 |
| DMEM | Gibco, Thermo Fisher Scientific | #21063-029 |
| Fetal bovine serum (FBS) | Gibco; Thermo Fisher Scientific | #10270-106 |
| Penicillin/Streptomycin | Gibco, Thermo Fisher Scientific | #15140-122 |
| Glutamax | Gibco, Thermo Fisher Scientific | #35050-038 |
| Poly-L-lysine (PLL). | Sigma-Aldrich | #P4707 |
| Neurobasal-A | Gibco, Thermo Fisher Scientific | #12348-017 |
| B27 | Gibco, Thermo Fisher Scientific | #17504-044 |
| Nocodazole | Sigma-Aldrich | #SML1665 |
| Cytochalasin-B | Sigma-Aldrich | #C6762 |
| Blebbistatin | Sigma-Aldrich | #B0560 |
| Laminin | Sigma-Aldrich | #P2020 |
| Lipofectamine 2000 | Invitrogen | #11668 |
| SPIONs (Fluid-MAG-ARA) | Chemicell | #4115 |
| Epoxy resin | Electron Microscopy Science | Epon 812 |
| **Commercial kits** |  |  |
| Macherey-Nagel NucleoBond Xtra Midi kit | Macherey-Nagel | #740412 |
| nucleoSpin RNA PLUS XS Kit | Macherey-Nagel, Düren | #740990.50 |
| Quant-iT RiboGreen RNA Kit | Thermo Fisher Scientific | #R11490 |
| PathScan® Acetylated Histone H3 Sandwich Elisa Kit | Cell Signaling Technology | #7232C |
| Piercetm BCA Protein Assay Kit | Thermo Fisher Scientific | #23225 |
| Diluent X | Cell Signaling Technology | #11083 |
| **Recombinant DNA** |  |  |
| pcDNA nesprin TS plasmid in DH5α cells | Addgene | #68127 |
| **Software and algorithms** |  |  |
| Fiji | ImageJ | [91] |
| NeuronJ | Fiji plugin | [95] |
| Gene Ontology analysis software | PANTHER | https://geneontology.org/ |
| RNAseq analysis software | Galaxy | https://usegalaxy.eu/ |
| GraphPad software, version 7.0 | GraphPad | https://www.graphpad.com/scientific-software/prism/ |
| Sample size | G power | G*Power for Windows |
| **Other** |  |  |
| Quad ibidi μ-Dish | Ibidi GmbH | #80416 |
| Microfluidic devices | XONA | #RD150 |
| Halbach-like cylinder magnetic applicator | Home-made | [96] |
| Diamond knife | Diatome | https://www.diatomeknives.com/knives/diamond_knife.aspx |
| Mesh copper grids | Electron Microscopy Science | G300Cu |

**Video S1.** 3D rendering of the nucleus of a control neuron

Movie_K1_24.06.20.avi

**Video S2.** 3D rendering of the nucleus of a stretched neuron

Movie_S1_24.06.20.avi

